# SHIP1 modulation and proteome characterization of microglia

**DOI:** 10.1101/2023.09.25.559310

**Authors:** Erpan Ahat, Zanyu Shi, Shaoyou Chu, Hai Hoang Bui, Emily R. Mason, Disha M. Soni, Kenneth D Roth, Michael James Chalmers, Adrian L Oblak, Jie Zhang, Jesus A Gutierrez, Timothy Richardson

## Abstract

Microglia, the resident macrophage in brain, has gained significant attention due to their involvement in neurodegenerative diseases. Disease associated microglia (DAM) have been identified at sites of amyloid-beta plaques and neurodegeneration. Understanding microglial states in the aging brain has become crucial, especially with the discovery of numerous Alzheimer’s disease (AD) risk and protective variants in genes such as *TREM2, CD33, APOE, ABCA7, PLCG2,* and *INPP5D*, which are essential to microglia function^1^. Here we present a thorough examination of microglia-like cell lines and primary mouse microglia at the proteomic and transcriptomic levels to help illuminate the roles these genes and the proteins they encode play in various cell states. This analysis serves as a guide to the exploration of potential therapeutic targets in the context of neurodegeneration. INPP5D, which encodes the SHIP1 protein, is essential for microglia function. SHIP1 has emerged as a target of interest having been nominated as a therapeutic target by three teams within the Accelerating Medicines Partnership for Alzheimer’s Disease (AMP-AD)^2^. In this study, we compared the proteomic profiles of wildtype, SHIP1 heterozygous knockout, and homozygous knockout primary microglia. Our findings revealed significant proteomic alterations only in the homozygous knockout of the SHIP1 gene, revealing its impact on the microglial proteome. Additionally, we compared the proteomic and transcriptomic profiles of BV2 and HMC3 cells with primary mouse microglia because these cell lines are often used as microglial cellular models. Our results demonstrated a substantial similarity between the proteome of BV2 cells and mouse primary cells, while notable differences were observed between BV2 and human HMC3 cells, with some shared characteristics. Since SHIP1 functions as a lipid phosphatase that modulates phosphatidylinositol (PI) species, we conducted lipidomic analysis to quantify different phosphatidylinositols (PIs), phosphatidylinositol monophosphate (PIPs), and polyphosphoinositides (PPIs) in the HMC3 and BV2 cells. Under basal conditions, PI(3,4,5)P3 and PI(3,4)P2 species were detected at extremely low levels, making confident quantification challenging; however, PIP species within the overall pool were significantly changed upon SHIP1 overexpression in HMC3. This in-depth proteomic analysis of both mouse and human microglia, complemented by targeted lipidomic studies, enhances our understanding of these cellular models. The similarities between primary mouse microglia and the BV2 cell line is especially encouraging, supporting the use of this model for further investigations into the role that SHIP1 and other potential drug targets may play in the regulation of microglial states.

## Introduction

Alzheimer’s disease (AD) is a devastating neurodegenerative disorder characterized by progressive cognitive decline and dementia. Extensive research has focused on understanding the pathological mechanisms underlying the abnormal accumulation of amyloid-beta plaques and neuronal neurofibrillary tangles, the hallmark features of AD^3^. Alongside these efforts, the study of dysregulated biological pathways underlying the disease process remains an area of intense research. Novel therapeutic targets within these pathways that can significantly alter the clinical course of AD require preclinical validation in models of this disease^4^.

Microglia, as the resident macrophage in brain, play a crucial role in regulating immune responses within the central nervous system (CNS)^5^. They are among the first responders to CNS injury and are vital for defending the neuronal microenvironment against infection, inflammation, and neurodegeneration^6^. Comprehensive single-cell RNA sequencing (RNA-seq) and analysis of AD and aging brain have identified numerous microglial states that are now being investigated in various normal and disease processes, including disease-associated microglia (DAM), that respond to neuronal damage^7^. However, dysregulated microglia triggered by environmental toxins, endogenous proteins, neuronal death, or infectious agents can lead to the production of inflammatory mediators that contribute to neuronal injury^8^.

Receptors expressed on microglia, such as triggering receptor expressed on myeloid cells 2 (TREM2), are known to play a role in the clearance of amyloid plaques and dead neuronal cells^9^. The phagocytic removal of amyloid plaques and the clearance of cellular debris are essential functions of microglia in regulating the brain microenvironment. TREM2 regulates microglial function and the response to amyloid and tau proteins as well as plaques and tangles they eventually form in Alzheimer’s disease. The lipid phosphatase SHIP1, a key protein downstream from TREM2, has been described as a negative regulator of signaling pathways. SHIP1 binds receptor immunoreceptor tyrosine-based inhibitory and activation motifs (ITIMs and ITAMs) where it competes with kinases, and converts phosphatidylinositol (3,4,5)-trisphosphate [PI(3,4,5)P_3_] to phosphatidylinositol (3,4)-bisphosphate [PI(3,4)P_2_].

Therefore, its inhibition or downregulation has been proposed as a strategy to increase the neuroprotective functions of microglia^10^. Since Microglia play critical roles in the regulation of the neuronal microenvironment and the progression of AD, mouse primary microglia and cell lines like BV2 and HMC3 cells are commonly used as cellular models for mechanistic analyses, compound screening, and lead optimization assays. However, it remains unclear whether results obtained from one cell model are translatable to another.

To gain a better understanding of the role of SHIP1 in microglia function and to explore the differences among various microglia models, we conducted a comprehensive proteomic characterization of microglia upon SHIP1 knockout or overexpression. Additionally, we performed a comparative analysis of the proteomic profiles of primary microglia, BV2 cells, and HMC3 cells as well as transcriptomic profiles of BV2 and HMC3 cells to identify similarities and differences in the expression of numerous critical pathways. Furthermore, we quantified different phosphorylated phosphatidylinositol phosphate (PIP) species through lipidomic analysis in human and mouse microglia following SHIP1 manipulation. This study provides a comprehensive dataset and knowledge base regarding the proteome and transcriptome of various microglia models, sheds light on the global proteomic changes induced by SHIP1 modulation and elucidates alterations in PIP species levels resulting from SHIP1 overexpression.

## Results

### SHIP1 knockout in primary microglia have distinct proteome compared to wildtype primary cells

SHIP1 is a phosphatidylinositol phosphatase that plays a key role in the regulation of signaling downstream from TREM2 and in turn microglia function. One key hypothesis is that genetic depletion or pharmacological inhibition of SHIP1 enhances the protective function of microglia and reduces the risk of disease progression in AD patients. Much effort has been given in developing inhibitors as well as SHIP1 knockout (KO) mouse models which became key tools to study SHIP1 function.

To determine the global effect of SHIP1 depletion, we performed a label free global proteomics comparison of WT, SHIP1 heterogeneous KO and homozygous KO mouse primary cells, which were isolated from the cortical tissue of neonate mice as previously described^11^ (Fig.1A; Supplemental Table 1A). We compared the proteomes of heterozygous KO to WT and homozygous KO to WT, and homozygous KO to heterozygous KO cells. With a cutoff point of fold change (FC)>=2 and adjusted P value of <0.01, we identified that there were total of 492 differentially expressed proteins with fairly even distribution at upregulation and downregulation in homozygous KO compared to WT (Fig.1A-B; Supplemental Table 1A). We performed a gene ontology (GO) analysis using Metascape to identify significantly altered pathways, we identified that Ribonucleoprotein complex biogenesis, mRNA processing, Golgi vesicle transport, cell division, and protein transport to plasma membrane function are among the top enriched pathways that differ in homozygous KO vs. WT phenotypes (Fig. 1C). This makes sense considering the importance of SHIP1 in the regulation of TREM2 dependent endocytosis, phosphoribosyl signaling and lysosome function^12^. Interestingly, the proteome of heterozygous knockout cells showed high similarity to that of WT (Fig.1D; Supplemental Table 1B). Only 17 proteins were identified to be significantly altered in SHIP1 heterogeneous KO compared to WT. The only significantly altered pathway in heterozygous knockout is the regulation of cytokine production (Fig.1E).

**Figure 1.**
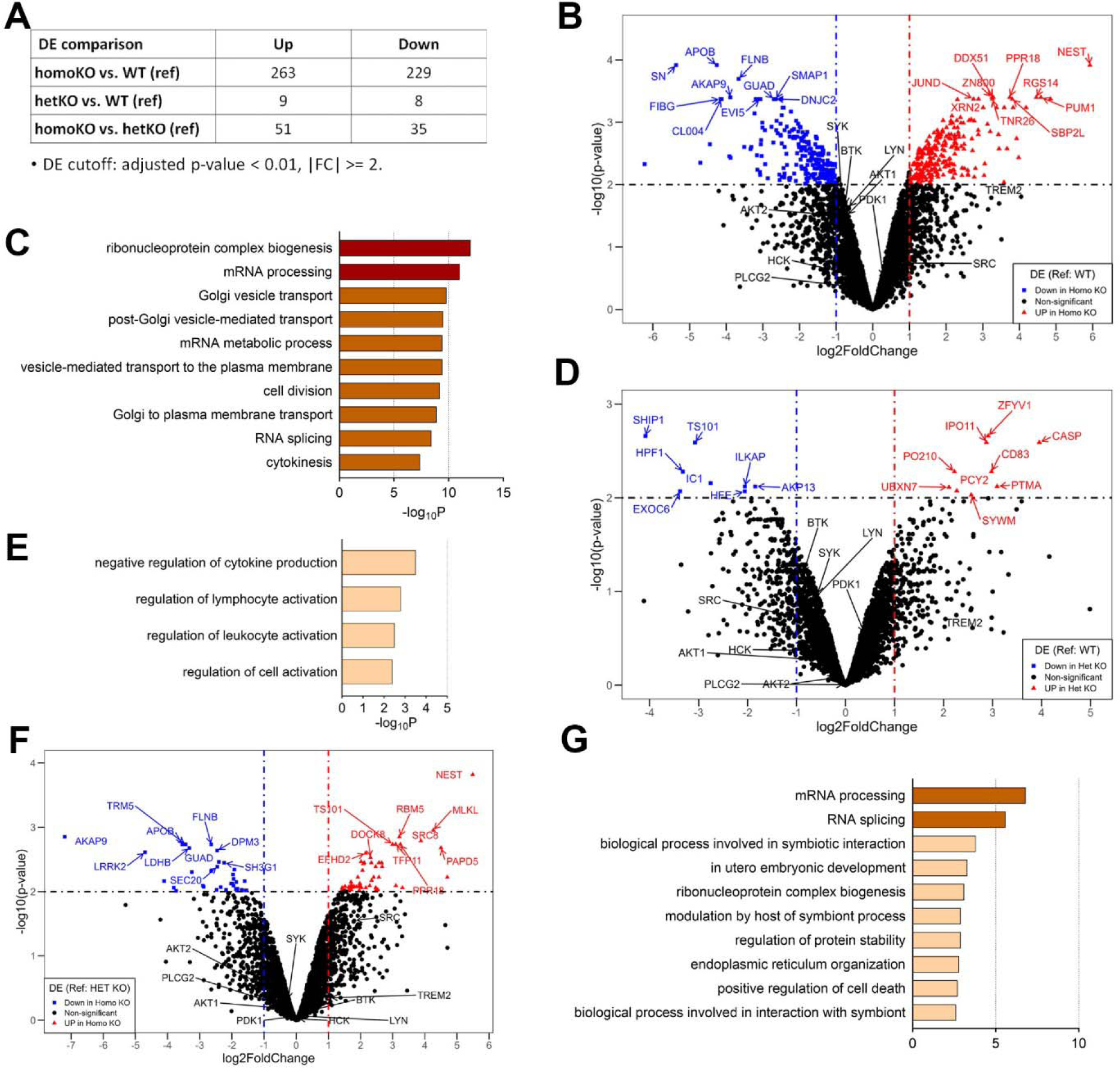
Comparative proteomics of WT vs. SHIP1 heterozygous knockout vs. SHIP1 homozygous knockout B6 mice derived primary microglia. **(A)** Summary table of upregulated (Up) and downregulated (Down) proteins in knockout cells. **(B)** Volcano plot of candidate proteins that are differentially expressed in homozygous knockout cells compared to WT. Significantly altered proteins are highlighted. **(C)** Metascape Gene Ontology (GO) enrichment analysis of significantly changed proteins in SHIP1 homozygous knockout proteins compared to WT. **(D)** Volcano plot of candidate proteins that are differentially expressed in heterozygous knockout cells compared to WT. Significantly altered proteins are highlighted. **(E)** Metascape Gene Ontology (GO) enrichment analysis of significantly changed proteins in SHIP1 heterozygous knockout proteins compared to WT. **(F)** Volcano plot of candidate proteins that are differentially expressed in homozygous knockout cells compared to heterozygous knockout cells. Significantly altered proteins are highlighted. **(G)** Metascape Gene Ontology (GO) enrichment analysis of significantly changed proteins in SHIP1 homozygous knockout compared to heterozygous knockout proteins.

Lastly, we also compared the proteome of homozygous knockout to that of heterozygous knockout cells (Fig.1F; Supplemental Table 1C). There were 86 significantly altered proteins and the top enriched GO terms were mRNA processing, symbiotic interaction, and regulation of protein stability among others (Fig.1G). This data shows the importance of using models of total SHIP1 knockout for the study of its function.

### Subcellular fractionation and comparative proteomics of primary microglia, BV2 and HMC3 cells

Even though primary microglia cell models are commonly used to study microglia biology, others including BV2 and HMC3 cells are widely used alternatives to primary cells because they are more readily available. There is no lack of discrepancies in the observations and conclusions reached when BV2 vs. HMC3 cells are used in studying TREM2 and SHIP1 biology^13, 14^. It is likely that the proteome differences in the cell lines account for the discrepancies. Here, we aimed to understand the similarities and differences of these cell models versus primary microglia at the proteome and transcriptome levels. We compared the proteomes of rodent BV2 cells to those of primary microglia (Fig. 2A-C; Supplemental Table 2A). The cells were sub-cellularly fractionated prior to proteomic analysis. As shown in Fig.2B for a membrane protein Lamp2, the fractionation procedure enriched Lamp2 mainly in the membrane fraction. TREM2 was also mainly enriched in the membrane fraction with some enrichment in the nuclear soluble fraction. We found that the proteome of BV2 and mouse primary microglia are largely similar with some differences (Fig. 2C). When the proteome of BV2 vs. HMC3 were compared, however, we found that there are significant differences in their respective proteomes, and multiple pathways were significantly different between the two including neurological diseases (Fig. 2D; Supplemental Table 2B). It is obvious from the data that BV2 cells have more similarity to that of primary mouse microglia than human HMC3 cells. One caveat on this conclusion is that, to compare the proteome of human cell line HMC3 to a murine cell line BV2, the mass spectrometric data for both were searched against a human database in which case only the common peptides (identical amino acid sequences) between BV2 and HMC3 were used for proteome comparisons. The reason for limiting analyses to homologous common peptides is to prevent biases that may occur due to differences in detection efficiencies between human and rodent peptide sequences. It is inherently unfair to compare the proteomics of two different species, but the above method is simple and actionable with stated limitations.

**Figure 2.**
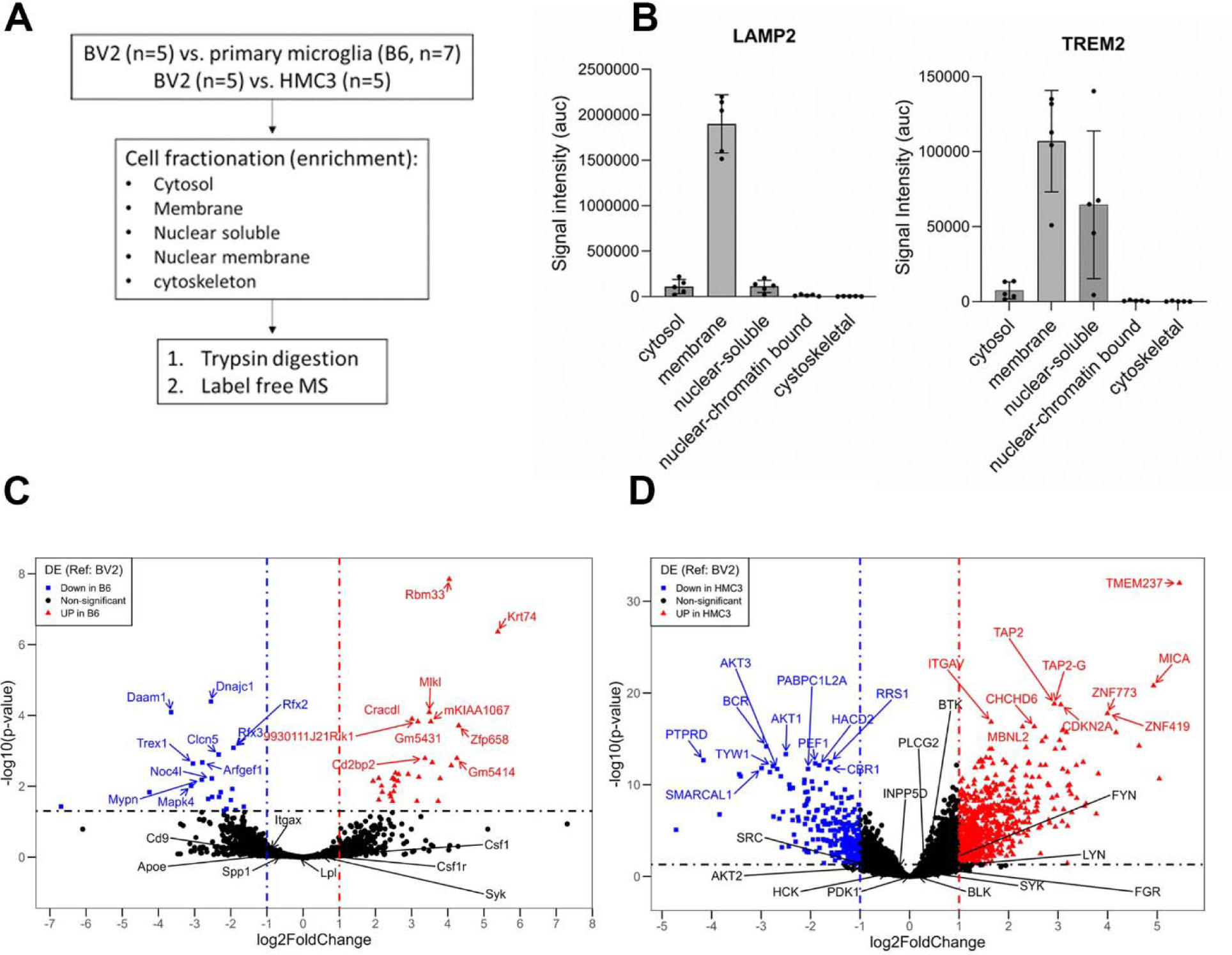
Comparative proteomics of primary microglia (B6) vs. BV2 vs. HMC3 cells. **(A)** Schematic of cell fractionation and sample preparation workflow. **(B)** Signal intensity of LAMP2, a lysosomal transmembrane protein, and TREM2 are shown in different fractions of BV2 cells. Note that LAMP2 signal mainly is present in the membrane fraction. **(C)** Volcano plot of proteins in BV2 and B6 proteome. Significantly changed proteins with fold change (FC) >=2 and adjusted P value of <0.05 are highlighted. **(D)** Volcano plot of proteins in BV2 and HMC3. Significantly changed proteins with fold change (FC) >=2 and adjusted P value of <0.05 are highlighted.

Considering that TREM2 is important to microglial function, we also assessed the expression of downstream proteins and other receptors (TLR4, FCGR2B, and CSF1R) in these cells, summarized in Table 1. Interestingly, we identified TREM2 in BV2 cells but not in HMC3 and primary microglial cells. SHIP1 protein levels were comparable between cell types while AKT1 levels were significantly higher in BV2 compared to primary microglia and HMC3 cells. TLR4 was significantly higher in HMC3 than BV2 which highlights differences between the cell types (Table 1).

**Table 1.** Selected proteins involved in microglial function and their differential expression in HMC3 vs. BV2 and B6 vs. BV2 cells.

| **Gene/protein** | **Quantified in**<br>**combined dataset:**<br>**<u>log2FC(HMC3/BV2)</u>** | **Quantified in**<br>**combined dataset:**<br>**<u>log2FC(B6/BV2)</u>** |
| --- | --- | --- |
| AKT1 | -2.5 | -1.7 |
| LYN | -0.77 | 0.6 |
| SHIP1 | -0.2 | 0.26 |
| PLCG2 | 0.29 | -9.5 |
| BTK | 0.36 | 0.09 |
| SYK | 0.36 | 0.44 |
| MTOR | 0.54 | 0.5 |
| RPTOR | 1.58 | -0.72 |
| TLR4 | 1.86 | -0.87 |
| APOE | NA | -0.71 |
| FCGR2B | NA | -1 |
| CSF1R | NA | 0.57 |

### Transcriptome and proteome comparison of HMC3 and BV2 cells

Considering that there is quite significant difference in the proteome of HMC3 and BV2 cells, and the inherent limitations of making proteome comparisons of two different species, we also performed transcriptome analysis. Comparing the transcriptome of HMC3 to that of BV2 with a cut off of absolute log_2_FC>=2 with a P<0.01, we identified numerous genes whose expression is significantly different between the two cell lines (Fig. 3A). GO term analysis showed that major differences are in pathways including regulation of cell activation, extracellular matrix organization and cell adhesion among others (Fig. 3B). The differences in these pathways could lead to differences in endocytosis and activation in BV2 and HMC3. We also searched for differences and similarities in transcripts whose corresponding proteins were present in our proteome data. Our results show consistency between proteome and transcriptome for gene products involved in fiber organization, signaling, nervous system development and focal adhesion (Fig. 3C-D). Lastly, when we reanalyzed the proteomics dataset for the proteins with transcripts that were identified in transcriptomics dataset, we identified that pathways such as RAB GEFS, mitochondrial organization, and autophagy were consistent between them. More importantly, our results show significant differences in the transcript level between HMC3 vs. BV2 (Fig. 3C-D) do not correlate well with less significant differences in the proteome of same genes for HMC3 vs. BV2 (Fig. 3E-F). This indicates that one has to be careful not to extrapolate too much from transcript level comparisons to proteome level changes in these cell lines.

**Figure 3.**
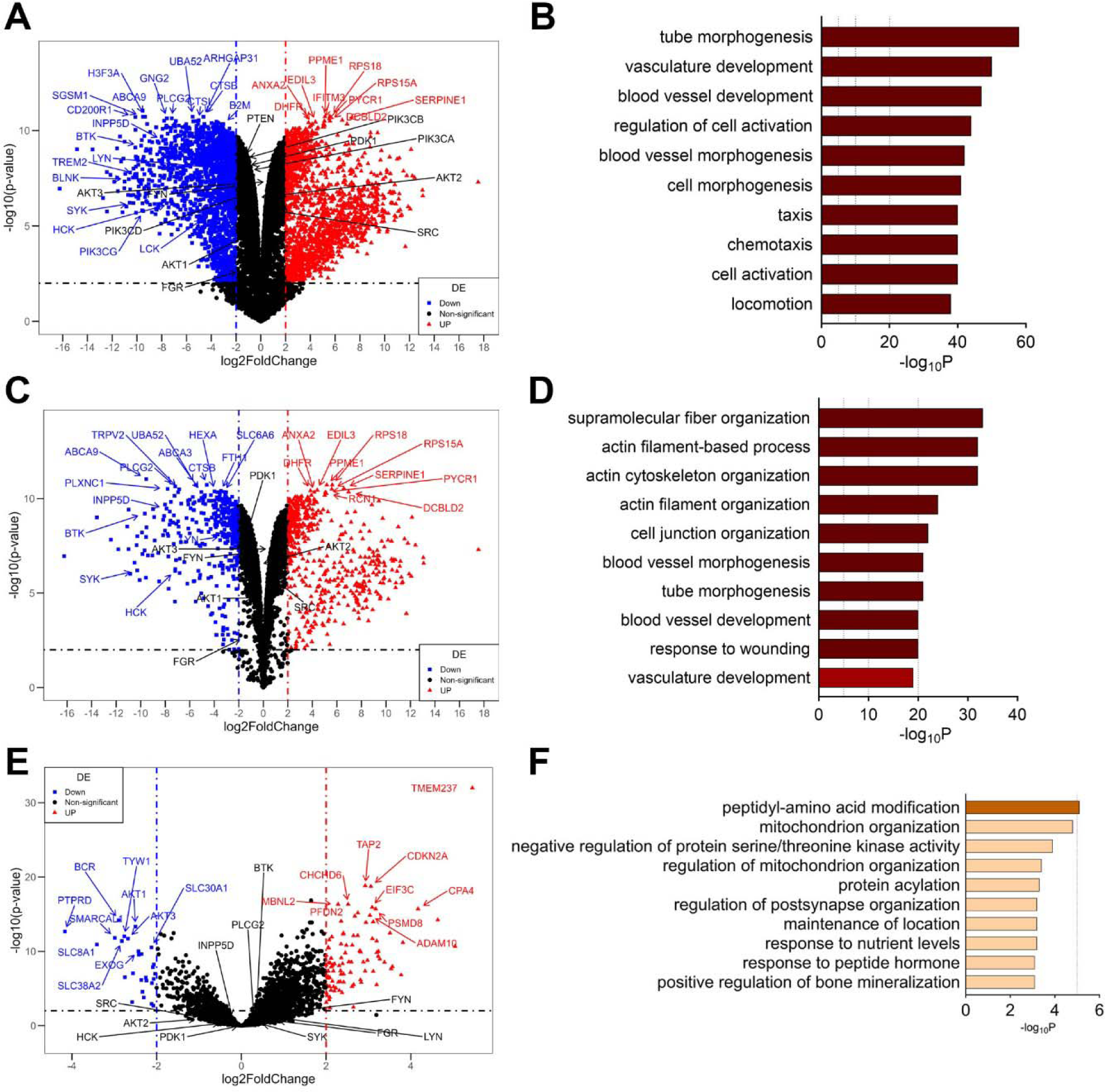
Transcriptomics and proteomics comparison of HMC3 and BV2 cells. **(A)** Volcano plot for HMC3 and BV2 transcript expression profiles. Significantly different genes with fold change log_2_FC >=2 and adjusted P value of <0.01 are highlighted. **(B)** Metascape Gene Ontology (GO) enrichment analysis of significantly altered genes in HMC3 compared to BV2 cells. **(C)** Volcano plot of genes in HMC3 and BV2 transcriptomics that overlaps with the HMC3 and BV2 proteomics data. Significantly different genes between the two cell lines with fold change log_2_FC >=2 and adjusted P value of <0.01 are highlighted. **(D)** Metascape Gene Ontology (GO) enrichment analysis of significantly altered genes that overlap with proteomics in HMC3 compared to BV2 cells. **(E)** Volcano plot of proteins in BV2 and HMC3 proteome that overlap with the corresponding genes identified in the transcriptomics data. Significantly changed proteins with fold change log_2_FC >=2 and adjusted P value of <0.01 are highlighted. **(F)** Metascape Gene Ontology (GO) enrichment analysis of significantly altered proteins that overlap with transcriptomics in HMC3 compared to BV2 cells.

### *Modulation of* SHIP1 *activity* regulates phosphatidylinositol (PI) and phosphatidylinositol phosphate (PIP) species levels

SHIP1 is a lipid phosphatase that dephosphorylates PI(3,4,5)P3 into PI(3,4)P2 to regulate TREM2 signaling and its downstream signaling cascade. With the notion that SHIP1 expression changes modulate PI(3,4,5)P3 and PI(3,4)P2 levels, and potentially other PIP species, we systematically quantified the levels of different PI and PIP species in two different microglia-like cell lines. WT and SHIP1 overexpressing HMC3 cells were harvested, and intracellular levels for various PIP species were quantified (Fig.4A). We identified that SHIP1 stable overexpression increases the level of PI but decreases the levels of different PIP species including PI(3)P, PI(4)P and PI(4,5)P2. Surprisingly, we did not detect any PI(3,4)P2 or PI(3,4,5)P3 in extracts from both cell lines. These data highlight the low levels of SHIP1 substrates PI(3,4)P2 and PI(3,4,5)P3 in these cells under our study conditions. The results for the other phosphoinositol species do demonstrate the effect of SHIP1 protein overexpression on total phosphoinositol and total PI(4)P levels, showing the dramatic remodeling of the phosphoinositol landscape in cells (Fig. 4A). BV2 cells are commonly used to screen and evaluate SHIP1 inhibitors. We also performed targeted lipidomic to quantitate the levels of various PIP species in BV2 cells. Various PIP species were detected in BV2 cells, but like HMC3 cells, the level of PI(3,4,5)P3 is extremely low and no PI(3,4)P2 was detected (Fig.4B).

**Figure 4.**
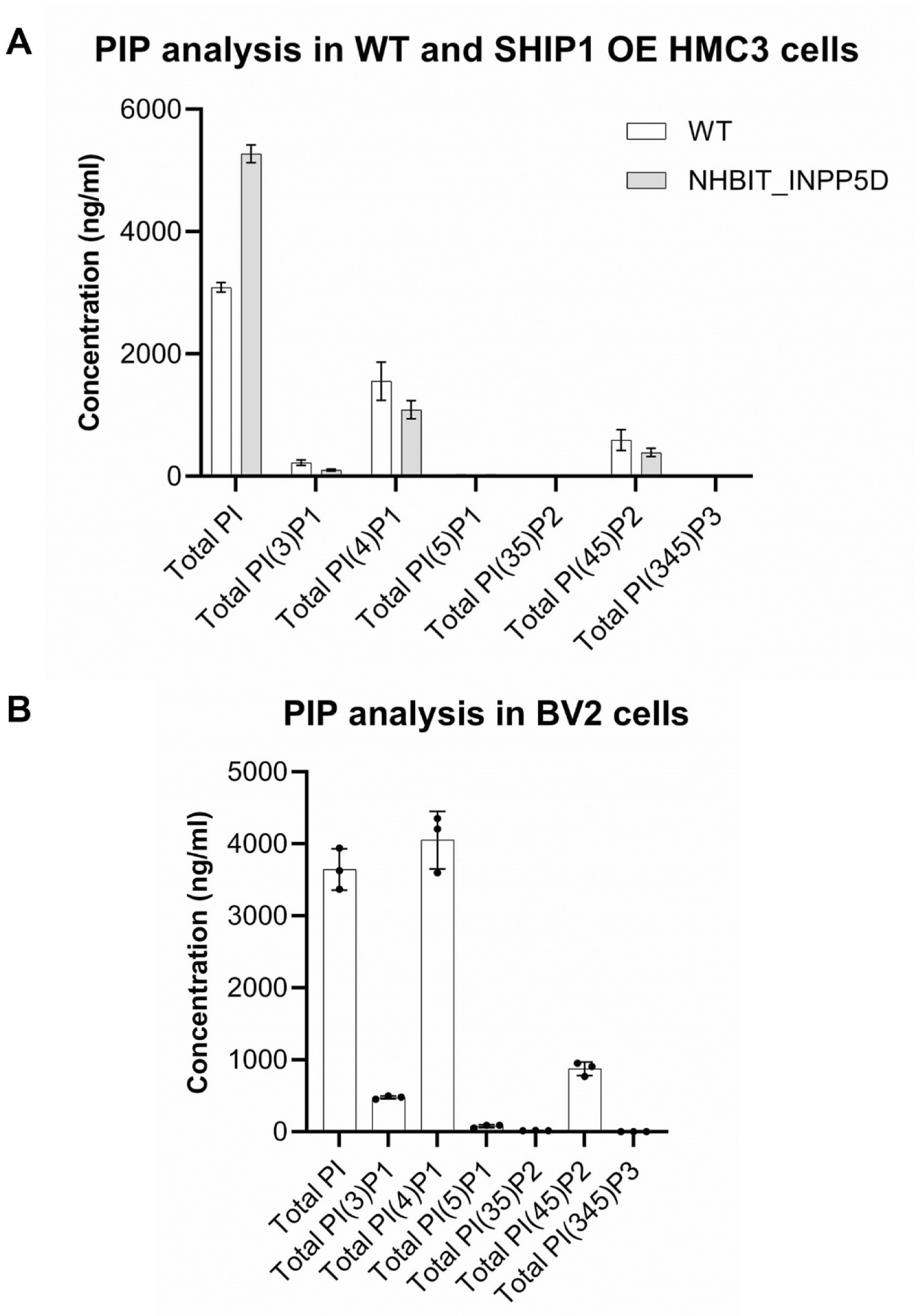
Quantification of different Phosphatidylinositol (PIP) species in BV2 and HMC3 cells. **(A)** Quantification summary of PIP species in wildtype (WT) and SHIP1 overexpressing (NHIBIT_INPP5D) cells. SHIP1 overexpression increased the level of PI but reduced the levels of all different phosphorylated species including PI(3)P, PI(4)P, PI(4,5)P2. Levels of PI(3,4,5)P3 levels were near background noise level and no PI(3,4)P2 were detected in HMC3 cells. **(B)** Quantification summary of PIP species in BV2 cells. Note that no PI(3,4)P2 species were detected in BV2 cells and PI(3,4,5)P3 levels were at noise levels.

## Discussion

Microglia, which are the key regulators of immune responses in the central nervous system, have emerged as crucial players in AD pathogenesis. The current study compared the proteomic profiles of primary microglia, BV2 cells, and HMC3 cells, which are commonly used as in vitro models of microglia. The analysis revealed that BV2 cells exhibited a greater similarity to primary microglia than HMC3 cells. Several pathways, including those associated with neurological diseases, showed significant differences between BV2 and HMC3 cells.

Considering that TREM2 signaling is a key focus in AD research, the study also assessed the expression of TREM2 and downstream proteins in these cell types. TREM2 was detected in BV2 cells but not in HMC3 or B6 primary microglial cells, indicating potential variations in TREM2 expression across different microglia models. SHIP1 protein levels were comparable between the cell types, while AKT1 levels were significantly higher in BV2 cells.

Since the role of SHIP1 in microglia function is of particular interest, proteomic analysis was performed on primary microglia with SHIP1 knockout, revealing distinct protein expression patterns compared to wild-type cells. The identified differentially expressed proteins were enriched in pathways related to Golgi vesicle transport, endocytosis, protein stability regulation, and lysosome function, which are all important processes involved in microglia activation and immune response. These findings highlight the significance of SHIP1 in the regulation of TREM2-dependent endocytosis and downstream signaling cascades^15^.

In addition to proteomic analysis, this study investigated the effects of SHIP1 modulation on phosphatidylinositol (PI) and phosphatidylinositol phosphate (PIP) species levels. SHIP1 overexpression in HMC3 cells resulted in an increase in PI levels and a decrease in various PIP species. However, PI(3,4)P2 and PI(3,4,5)P3 species were not detected, suggesting that longer-term effects may be necessary for observing changes in these specific PIP species. Similarly in BV2 cells, the levels of PI(3,4)P2 and PI(3,4,5)P3 were undetectable with our methods. These findings suggest that stimulating the PIP pathway to increase the levels of PI(3,4)P2 and PI(3,4,5)P3 prior to SHIP1 modulation may be necessary to evaluate the direct effects of SHIP1 overexpression on PIP species levels.

The comprehensive proteomic characterization of microglia and the comparison of different cell models provide valuable insights into the role of SHIP1 in microglia function and its potential as a therapeutic target for AD. The results demonstrate that SHIP1 knockout leads to distinct proteomic changes in primary microglia, emphasizing its importance in regulating various pathways associated with microglia activation. Furthermore, the differences observed between BV2 and HMC3 cells highlight the need for careful selection of microglia models in studying specific aspects of AD pathogenesis, particularly the TREM2 pathway. In addition to BV2 and HMC3 cells, iPSC derived microglia are being used as a microglia model system^16, 17^.

Although this study provides important findings, there are certain limitations to consider. The study focused on proteomic and lipidomic changes and did not explore functional consequences or downstream effects of SHIP1 modulation in microglia cell physiology. Future studies should investigate the functional implications of the identified proteomic changes and explore the effects of SHIP1 manipulation on microglia-mediated neuroinflammation, phagocytosis, and clearance of amyloid plaques.

In conclusion, this study elucidated the proteome differences associated with SHIP1 knockout in primary microglia and highlighted the differences between various microglia cell models. The findings emphasize the importance of SHIP1 in regulating key pathways involved in microglia activation and immune response. Furthermore, the study provided insights into the effects of SHIP1 modulation on PI and PIP species levels, which have implications for understanding the downstream signaling events in microglia. These findings contribute to our understanding of microglia biology and provide a foundation for future studies investigating SHIP1 as a potential therapeutic target for AD.

## Materials and Methods

### Reagents

Dithiothreitol (Sigma, Cat# D0632; DTT); Iodoacetamide (Sigma, Cat# A3221; IAA); 1M TEAB, pH 7.5 (Thermo Scientific, Cat# 90114); Trypsin (Promega, Cat# V5280); S-Trap^TM^ mini-MS sample prep kit (Fisher Scientific, Cat# NC1828287); 100% Acetonitrile (Burdick & Jackson. Cat# 015-4).

### Cell culture, transfection, and treatment

For cell culture, BV2 and HMC3 cells were grown in DMEM (Invitrogen) containing 10% super calf serum (Cytiva, Cat# SH30898.03) and 1% Pen/Strep (Fisher Scientific, Cat#15140122) at 37°C in a 5% CO2 incubator. Primary microglia cells were collected from P0 pups. BV2 and HMC3 proteomics comparison was done at cell passage of P13 and P18 for BV2 and HMC3, respectively. PIP analysis was done at cell passage of P18 and P20 respectively for WT HMC3 and HiBit-INPP5D HMC3 cells, and P17 for BV2 cells.

### Subcellular fractionation

Subcellular fractionation was carried out following the manufacture’s protocol (MAN0011667_Subcellular_Protein_Fraction_CulturedCells_UG). Briefly, 500 μL of CEB was added to cell pellet and gently mixed at 4°C for 10 minutes. Samples were centrifuged at 500 × g for 5 minutes and the supernatant was collected into a clean pre-chilled tube as the cytoplasmic fraction. 500 μL of ice-cold MEB was applied to pellet, vortexed for 5 seconds and incubated at 4°C for 10 minutes with gentle mixing. Membrane fraction is collected by centrifugation at 3000 × g for 5 minutes. After that, 250 μL NEB containing protease inhibitors were added to pellet, vortexed for 15 seconds and incubated at 4°C for 30 minutes with gentle mixing. Soluble nuclear extract was collected by centrifugation at 5000 × g for 5 minutes and transferred to a fresh tube. Next, 250 μL NEB containing protease inhibitors, CaCl2 and Micrococcal Nuclease were added to pellet, vortexed for 15 seconds and incubated at room temperature for 15 minutes. After incubation, samples were vortexed for 15 seconds and centrifuged at 16,000 × g for 5 minutes, and the supernatant was collected in a fresh tube as the chromatin-bound nuclear extract. Finally, 250 μL PEB solution containing protease inhibitors was added to the pellet, vortexed for 15 seconds and incubated in room temperature for 10 minutes. Samples were centrifuged at 16,000 × g for 5 minutes and the supernatant was transferred to a new tube as the cytoskeletal fraction. Samples were dried under low vacuum prior to processed with S-trap proteomic protocol.

### S-trap based proteomic sample preparation

The S-trap procedure was performed following the manufacturer’s protocol for S-trap kit (S-Trap^TM^ mini-MS sample prep kit: 100 – 300 μg https://www.fishersci.com/shop/products/s-trap-minikit-10x100-300ug/NC1828287), briefly: cell pellets were lysed with 100 μL of 1x SDS buffer (final SDS is 5%) followed by addition of 5 μL of 100 mM DTT and incubation at 55 C for 15 min. 5 μL of 400 mM iodoacetamide was added to the mixture and incubated at room temperature in dark for 10 min. Samples were acidified with 10 μL of 12% phosphoric acid. Samples were then mixed with 500 μl 100 mM TEAB in 90% methanol and applied to S-Trap column, centrifuged at 10,000g for 30 sec to trap proteins followed by 3x wash with 400 μL of Buffer 5. Samples were digested with 0.1 μg/μL trypsin in 50 mM TEAB, pH 8 overnight at 37 °C. The digested samples were eluted sequentially with 80 μL of 50 mM TEAB, pH 8, 80 μL of 0.2% aqueous formic acid, and 80 μL of 50% acetonitrile in water. Samples were dried down under low vacuum (Genevac) at room temperature. The pellets were resuspended in 50 mM TEAB, pH 8 and stored at -80°C degree for further analysis using mass spectrometry.

### Liquid chromatography mass spectrometry and Data analysis LC Conditions

An Ultimate 3000 HPLC system was run at 300 nL/min. at 1% B with Solvent A as 0.1% formic acid in water (Thermofisher Optima LS118-4) and Solvent B as 0.1% formic acid in acetonitrile (Thermofisher Optima LS120-4). Ten microliters of each sample were loaded onto a trap column (100 mm i.d. x 2 cm; Acclaim PepMap C18, 3 mm, 100A; ThermoFisher 164564) in 0.1% formic acid in water. Analytical column was the Thermo pepmap RSLC (75 mm x 50 cm, C18, 3 mm, 100A; Thermofisher ES803A). Analytical column separation was at 300 nL/minute with gradient of solvent B consisting of 1 to 85% in 103 minutes followed by 4 minutes at 85% solvent B at 30°C. Michrom Bioresources bovine protein tryptic digest standards (lactoglobulin, lactoperoxidase, carbonic anhydrase, and glutamante dehydrogenase; catalogue numbers PN 60006, PN 60011, PN 60007, and PN 60010, respectively) were analyzed as instrument performance controls.

### Mass Spectrometer Conditions

A Fusion Lumos mass spectrometer was operated using data dependent acquisition mode with the following parameters: full scan (m/z range 350-1500, OT resolution 240000, intensity 2.5e4). Quadrupole isolation window was set at 1.2 m/z. Mass analysis was performed using HCD-MS2 in ion trap. MS/MS spectra were acquired by HCD of 28% and normal AGC target. Precursor ions selected for fragmentation were put on a dynamic exclusion list for 60 seconds.

### Mass spectrometry data analyses

Resulting mass spectrometric data files (.Raw) files were processed using standard Bottom-Up proteomic analyses with an internal proteomic data processing software suite previously described^18^. Briefly, peptides were identified using X!Tandem and Omssa peptide search engines with trypsin as the enzyme with a maximum of 2 missed cleavage sites, fixed carbamidomethyl modification at cysteines (+57.0215), and variable modifications for methionine oxidation (+15.9949), asparagine and glutamine deamidation (+0.9840), and phosphorylation (+79.9663) at serine, threonine and tyrosine residues. Variable n-terminal acetylation (+42.0106) was also allowed. False positive identifications were controlled by the reversed protein database analyses and false discovery rates (FDR; q-values) were estimated as described previously^18^. Chromatographic alignments were performed for each file and used to normalize peptide retention times and guide peptide quantifications (XIC; extracted ion chromatogram peptide integrations)^18^. XIC values of less than 2^15^ were considered background and removed prior to subsequent data analyses. Reported proteins with less than 2 distinct peptides were also removed from additional analyses.

### Phosphatidylinositol Phosphate (PIP) species quantification using mass spectrometry **Lipid Extraction and Sample Preparation.**

Phosphatidylinositols and phosphoinositides were extracted in cells using a modified Bligh & Dyer two-phase extraction that was previously used^19^. 12.5 μL of 10 μg/mL internal standard solution was added, followed by 400 μL of methanol and 300 μL of dichloromethane. Samples were then shaken on a tissuelyser (Qiagen, Valencia CA) for five minutes. Another 300 μL of dichloromethane and 300 μL of 1.0 N HCl were added to separate the phases. Sample were then shaken on a tissuelyser for another 5 minutes for extraction. After centrifuging the samples for 15 minutes at 5°C at 4000 rpm on bench top centrifuge 5810R (Eppendorf, USA), 600 μL of dichloromethane (bottom layer) was transferred to LCGC-certified clear glass vial. After drying down under a warm stream of nitrogen, the extract plasma was then reconstituted with 5 μM EDTA and 0.04% DiiPEA in 80% methanol.

### UHPLC Conditions

LC/MS/MS analysis of PIs, PIPs, and the PPIs was performed using a Sciex 7500 mass spectrometer interfaced with an ultra-high performance liquid chromatography (UHPLC) system in negative ESI mode. The UHPLC system consisted of an Agilent 1290 binary pump, thermostat, thermostat column compartment, and autosampler. The injection volume was set to 2 μL. Samples were chromatographically resolved using a Kinetex EVO C18 UHPLC column, 2.1x150 mm, 1.7 μm (Phenomenex, CA, PN:00F-4726-AN) held at 60°C. Mobile phase A was 5 μM EDTA and 0.04% DiiPEA in water. Mobile phase B was 5 μM EDTA and 0.04% DiiPEA in acetonitrile. This assay was broken into three different isocratic conditions that were used to separate all PIs, PIPs, and PPIs. First method, an isocratic LC condition was kept at 62% mobile phase B for 8 minutes to separate PI species. Second isocratic method, mobile phase B was kept at 38% for 10 minutes to analyze PIP species. Last, a shallow gradient method was used to separate PIP_2_ and PIP_3_ species. It used a gradient that gradually increased from 33% B to 35% B over 6 min, then the column re-equilibrated with 33% mobile phase B for 3 min. For each analytical run, all samples were analyzed using Method 1, followed immediately by Method 2 and finishing with Method 3. The advantage of this staged sample analysis is that it greatly reduces the total amount of re-equilibration time for the two isocratic method and all three methods can be set up to run overnight on the same system. For all conditions, the solvent flow rate was 0.40 mL/min. The valve, sample loop, and needle were washed with a solution of 5 μM EDTA in 50% methanol/50% acetonitrile for 30 sec.

### Calibration Standards and Quantification

Different PI, PIP and PPI species were quantified using standard curves based on the ratios of the peak areas of analytes to internal standards. The calibration curves of the closest counterpart were used for those PIs, PIPs, and PPIs where authentic standards were not available. Calibration curves were calculated by least-square linear regression with 1/x weighting. Data analysis was performed using Multiquant 3.0 (Sciex, Framingham, MA). The raw data were exported to Excel spreadsheets. JMP 15 was used for statistical analysis.

### Quantification and Statistical analysis

For transcriptomic and proteomic data analysis, all data preprocessing, differential expression analysis, and graph plotting were performed in R version 4.2.1.

For the proteomic data from cell line or primary microglia, first, data preprocessing was conducted for probe filtering and missing values imputation. Probes with more than 50% missing values in each group were removed, and for the rest of the probes, the mean of non-zero values were used to impute the missing values in each group. Next, quantile normalization was performed for the entire data to remove sample batch effects. In the case of the differential expression analysis, R limma package was used to perform the three set of comparisons (i.e., SHIP1 heterozygous KO to WT, SHIP1 homozygous KO to WT, and SHIP1 homozygous KO to SHIP1 heterozygous KO cells), and to compare among primary microglia (B6), BV2, and HMC3 cells. In the case of BV2 mouse microglia cell line and HMC3 human microglia cell line comparison, only ortholog proteins present in both species were used for comparison. In limma package, linear modeling approaches were implemented by lmFit combined with the empirical Bayes statistics applied by eBayes. P-values were further adjusted by Benjamini & Hochberg (BH) method and foldchange was calculated in top Table to extract top-ranked genes or proteins, which were selected by arbitrary thresholds of adjusted p-values and fold changes mentioned in the Results section. In addition, the R ggplot2 package was used to plot volcano plots and highlight significantly altered genes.

For the BV2 and HMC3 transcriptomic data analysis, the same data preprocessing strategies and differential expression analysis tool as above were applied on the TPM (Transcripts Per Kilobase Million) data of RNA-seq from BV2 and HMC3 cell lines. Since we are using the normalized counts data, the normalization step in limma was omitted. To crosscheck the differentially expressed genes and proteins in the HMC3 vs. BV2 (with BV2 as reference) comparisons, the top-ranked genes from the HMC3 vs. BV2 differential gene expression analysis (using a cutoff of |log_2_FC| >=2 with BH adjusted P<0.01) were further crossed-checked to obtain the overlapped genes with the corresponding proteins in the above proteomic comparison and the significantly different genes and proteins in both datasets were shown in corresponding tables. Volcano plots were plotted using ggplot2 package to highlight overlapped significantly different genes and proteins.

## Acknowledgments

We thank Richard Higgs for his valuable suggestions for proteomics data analysis.

## Author Contributions

E.A., M.J.C, S.Y.C, A.L.O, T.R and J.A.G. designed the experiments; A.L.O prepared the primary microglia; S.Y.C generated the wildtype and INPP5D clones of HMC3 cells; E.A. performed the sample preparation and proteomics analysis. A.L.O performed the transcriptomics analysis. Z.S and J.Z performed the statistical analysis of the proteomics and transcriptomics data. H.H.B and K.D.R performed the lipidomic analysis. All authors discussed the results and contributed to the final manuscript.

